# Long-read transcriptomics highlights venom gland specialization and Inhibitor Cystine Knot (ICK) rich toxin diversity in Philippine tarantulas

**DOI:** 10.64898/2026.07.14.737467

**Authors:** Lorenz Rhuel P. Ragasa, Maria Mikaela U. Dumbrique, Sid Alfred S. Gamboa, Anj Guillan M. Baile, Darrell C. Acuña, Heidie L. Frisco-Cabanos, Ricardo C. H. del Rosario, Leonardo A. Guevarra, Myla R. Santiago-Bautista

**Affiliations:** Research Center for Natural and Applied Sciences, University of Santo Tomas, Sampaloc, Manila 1008, Philippines; Department of Biological Sciences, College of Science, University of Santo Tomas, Sampaloc, Manila 1008, Philippines; Department of Biochemistry, Faculty of Pharmacy, University of Santo Tomas, Sampaloc, Manila 1008, Philippines; Graduate School, Polytechnic University of the Philippines, Sta. Mesa, Manila 1016, Philippines; Philippine Arachnological Society, Inc., Paco, Manila 1007, Philippines; DOST-PCHRD Balik Scientist Program; Broad Institute of MIT and Harvard, Cambridge, MA, USA

**Keywords:** Theraphosidae, Oxford Nanopore sequencing, Venomics, Comparative transcriptomics, Orthogroups, Venom evolution

## Abstract

Animal venoms are a rich source of bioactive molecules, yet their diversity remains incompletely characterized in many species. Here we present the first long-read transcriptomic analysis of venom glands from Philippine tarantulas (Theraphosidae), a highly endemic but understudied group. Using Oxford Nanopore sequencing, we reconstructed near full-length venom gland transcriptomes across multiple species and identified extensive repertoires of toxin-encoding peptides. Venom glands were enriched in cysteine-rich inhibitor cystine knot (ICK) peptides, which dominated the toxin landscape and are known modulators of ion channels. Cross-species comparative analyses revealed a distinct transcriptional signature separating venom from non-venom tissues, driven by coordinated expression of toxin-associated and regulatory gene families. Phylogenomic reconstruction based on orthologous peptides recovered expected taxonomic relationships while revealing potential lineage-specific diversification and potential cryptic taxa. Despite a conserved core set of toxin families, substantial variation in toxin composition was observed among species, consistent with rapid evolution driven by gene duplication and functional divergence. Analysis of highly expressed ICK peptides showed a conserved cysteine framework alongside marked sequence variability in inter-cysteine regions, supporting a model in which structural stability is maintained while functional diversification proceeds. Together, these findings establish the first long-read transcriptomic resource for Philippine theraphosid spiders, reveal a conserved molecular signature underlying venom gland specialization, and provide new insights into the diversification of ICK toxin repertoires that may facilitate future evolutionary and functional studies, including the discovery and characterization of bioactive venom peptides.

## Introduction

Venomous organisms have been a valuable source of bioactive compounds with significant therapeutic potential. Animal venoms are complex mixtures composed of peptides, proteins, and other small molecules that have evolved to target physiological pathways with high potency and selectivity [1,2]. Understanding the peptide diversity within these venoms is crucial in examining their evolutionary advantage, as well as potential biomedical uses. Among venomous arthropods, tarantulas (Family: Theraphosidae) produce diverse peptide toxins that primarily modulate ion channels, namely voltage-gated sodium (Na_v_), potassium (K_v_), and calcium (Ca_v_) channels [3,4]. These ion channel-modulating peptides have attracted considerable interest for their applications in pain management, neuropharmacology, antimicrobial development, and cancer therapeutics [2,5].

The Philippines, recognized as a global biodiversity hotspot with exceptionally high endemism [6], likely harbors unique, lineage-specific toxin repertoires in its fauna. However, despite the increasing global interest in spider venomics, no comprehensive characterization of venom peptides from Philippine tarantulas exists. Venom peptides evolve rapidly in response to prey interactions and ecological adaptation, often resulting in gene duplication, neofunctionalization, and lineage-specific toxin expansion [1]. Therefore, investigating venom peptide diversity in understudied lineages is essential to understand both toxin evolution and potential biomedical applications.

Advances in high-throughput sequencing, particularly in transcriptomics, have transformed venom research by large-scale identification of toxin-coding genes directly from venom glands [7]. One currently popular sequencing technology is long-read sequencing, which, compared to short read sequencing, long-read technology can reconstruct complete transcripts, thereby resolving isoforms, and accurately delineating full-length transcript precursors, including signal peptides, propeptide regions, and mature toxin domains [8,9]. Thus, venom gland long read transcriptomics offers a useful framework for examining toxin precursor architecture, expression patterns, and venom gland specialization.

Because the molecular architecture and comparative diversity of venom-associated transcripts in Philippine tarantulas remain poorly resolved, there is still a limited basis for evaluating toxin repertoire variation across taxa. Moreover, although venom gland transcriptomes have been reported for several spider species, these studies have largely relied on short-read sequencing or have focused on non-Philippine taxa, leaving a substantial gap in our understanding of the full-length venom gland transcriptomes of Philippine theraphosids. Hence, this study aims to generate venom gland transcriptomes using long-read sequencing technology together with *de novo* assembly to reconstruct near full-length transcripts. High-quality transcripts were filtered and annotated to identify putative toxin families. Orthologous venom peptides were used to infer phylogenetic relationships among the studied taxa. Phylogenetic reconstruction based on venom gene repertoires provides insight into toxin diversification, species divergence, and potential cryptic lineages [9–11].

Importantly, this study represents the first long-read venom gland transcriptomic analysis for the Theraphosidae family and provides novel insights into toxin precursor diversity within the understudied theraphosid lineage. Integrating full-length transcripts with comparative venom peptide profiling and phylogenetic analysis across three genera highlights interspecific variability and suggests the presence of putatively new taxa. These findings establish a foundational molecular framework for future functional, evolutionary, and biomedical investigations of tarantula venoms.

## Materials and Methods

### Sampling, Species identification, and Animal care

A total of twelve live tarantula specimens were collected from selected localities in the Philippines with permits provided by the Department of Environment and Natural Resources – Biodiversity Management Bureau (DENR-BMB) (Gratuitous Permit No. 318). Sample metadata, including sample ID, collection coordinates, and species designation, were curated and standardized. Specimens were maintained individually under controlled laboratory conditions prior to tissue sampling. General husbandry followed standard practices for theraphosid spiders, including housing in individual containers under ambient temperature conditions and provision of water *ad libitum*.

Feeding was conducted at regular intervals, once every 2 weeks. To minimize potential contamination of venom gland transcriptomes by recently ingested prey material, individuals were not fed for a sufficient period, minimum of 3 to 7 days prior to dissection. Specimens were monitored to ensure normal behavior and condition prior to tissue collection.

### RNA extraction

Dissecting materials were treated with RNase decontamination solution, washed with sterile nuclease free water, then sterilized by autoclave. An hour before dissection, the individual was placed in a 4°C refrigerator for euthanasia, while ensuring the tissue will not be frozen. The individual was placed under a scanning stereomicroscope for dissection. For venom gland extraction, both of the chelicerae were removed from the body, then incisions were made to reveal the venom glands. Two whole intact venom glands were carefully removed and placed in a microcentrifuge tube for further processing. For small specimens, both whole chelicerae, which contains the venom glands, were used. Non-venom tissues, such as the remaining body, leg, and abdomen, were also extracted.

The tubes containing the tissues were spinned down and immediately immersed in liquid nitrogen. While frozen, the tissues were homogenized using a micropestle and subjected to RNA extraction using the Qiagen RNeasy^®^ Mini Kit (Cat. No. 74104), following manufacturer’s protocol with modifications. Tissues were lysed using 350 μL of Buffer RLT and were further homogenized by pipetting using a 10-μL tip. For the elution step, 30 μL of RNAse free water was used. RNA quantification and quality check were performed using a Denovix QFX fluorometer and Agilent TapeStation system. All steps, including centrifugation (12,000 × g), were performed at room temperature.

### Transcriptome sequencing, assembly and annotation

Full-length cDNA libraries were prepared from total RNA following the Oxford Nanopore Technologies (ONT) cDNA-PCR sequencing protocol, according to the manufacturer’s instructions (ONT cDNA-PCR Sequencing, SQK-PCS111). Library preparation was performed using ONT-recommended reagents, with bead-based cleanup steps carried out using Magbio HighPrep RNA elite and HighPrep PCR magnetic bead systems. The library was loaded into an R9.4.1 flow cell (FLO-MIN106D). Sequencing was performed by a Mk1B device using MinKNOW 24.11.10, with sufficient read coverage targeted for downstream transcriptome assembly and comparative analyses. The raw pod5 files were subjected to basecalling using guppy v6.5.7 in Super Accurate mode (SUP) with model dna_r9.4.1_450bps_sup. Reads were trimmed, oriented, and filtered using Pychopper v2, using the pHMM model to identify and remove adapter sequences.

The high-quality transcriptome assembly was created using RNABloom v2.0.1 [12], with a minimum long-read read depth of 20 (-lrrd 20) and two iterations of read error-correction (-errcorritr 2). This read-depth threshold was applied to reduce the influence of low-support transcripts and noisy abundance estimates in downstream expression analyses. Transcript abundance was estimated using NanoCount v1.0.0 [13], while open reading frames and peptide sequences were predicted using TransDecoder v5.5.0, retaining BLASTp [14] evidence. Peptides of at least 50 amino acids in length were included to allow recovery of small toxins.

To obtain comprehensive functional annotations of the predicted peptide sequences, multiple annotation databases were used. Sequence similarity searches were performed against UniProt and curated toxin databases such as ArachnoServer [15], the KNOTTIN database [16], and the animal toxin annotation project (Tox-Prot) dataset of UniProt [17], using DIAMOND v0.9.19 [18]. Functional annotations including Gene Ontology (GO) terms were retrieved from eggNOG-mapper [19,20] via Galaxy v2.1.13+galaxy0 [21] and UniProt-derived annotations. Additional feature predictions, including signal peptides via SignalP 6.0 [22], propeptide cleavage via ProP 1.0 [23], and toxin-associated properties via ToxinPred 3.0 [24], were incorporated. All annotation sources were integrated into a consolidated annotation table at the peptide-level, which served as the basis for downstream functional, toxin classification, and enrichment analyses.

### Toxin identification and comparative analysis

To enable comparative analysis across samples and species, orthologous relationships among predicted peptides were inferred using OrthoFinder v3.1.0 [25]. Peptides were clustered into orthologous groups (orthogroups) that represent putative homologous families, allowing the identification of shared, lineage-specific, and tissue-associated peptide groups. The overlap and distribution of orthogroups across samples were visualized using UpSetR in R [26]. Furthermore, a species tree inferred by OrthoFinder was included to provide phylogenetic context for these comparisons.

Transcriptomes obtained from venom glands were compared with those from non-venom tissues to evaluate venom gland-associated expression patterns of toxin transcripts. Because analyses were performed across multiple species without a shared reference, expression comparisons were conducted at the orthogroup level and interpreted within a relative framework rather than as formal differential expression. The transcript abundance values of each sample, measured in Transcripts Per Million (TPM), were aggregated at the orthogroup level by summing expression across all transcripts assigned to each orthogroup, reflecting the cumulative transcriptional output of gene families. Orthogroup expression values were then log_2_-transformed (log_2_(TPM + 1)). To reduce noise from low-variance orthogroups and focus the analysis on orthogroups contributing most strongly to expression differences among samples, the top 20% most variable orthogroups (n=1908 of 9521) were retained for principal component analysis (PCA). To assess the robustness of the resulting clustering patterns, PCA was repeated using alternative variance thresholds (top 10%, 30%, and 40% most variable orthogroups), which produced qualitatively similar sample groupings (Supplementary Fig. 1).

To interpret the primary axis of variation, orthogroups were ranked based on their loadings along PC1. Gene Ontology (GO) annotations from the consolidated annotation table were mapped to orthogroups by assigning each orthogroup all GO terms associated with its member peptides. Gene Set Enrichment Analysis (GSEA) was performed using clusterProfiler [27], with orthogroups ranked according to their PC1 loadings. The ten most significant GO terms were visualized as a bar plot showing normalized enrichment scores (NES) direction and magnitude. To further examine ontology-specific patterns, separate GSEA analyses were then performed for the Biological Process (BP) and Molecular Function (MF) ontologies. These results were visualized using clusterProfiler dot plots showing gene ratio, adjusted p-value, and gene set size. P-values were adjusted for multiple testing using the Benjamini–Hochberg method.

In parallel with orthogroup-based analyses, toxin-specific identification and classification were performed to characterize venom-associated transcripts. A multi-step annotation-based filtering strategy was employed to capture peptide toxins while minimizing false positives. Strong toxin-specific evidence was defined based on sequence similarity to curated toxin resources, including ArachnoServer, the KNOTTIN database, and the UniProt Tox-Prot dataset. Matches with E-values ≤ 1 × 10^-5^ were considered significant. Additional toxin-associated evidence included the presence of the Gene Ontology term “toxin activity” (GO:0090729).

In addition to sequence similarity evidence, peptides were further evaluated based on characteristic features of spider venom peptides, including the presence of signal peptides using SignalP, predicted toxicity using ToxinPred, and cysteine-rich frameworks. We further incorporated supplementary evidence from toxin-related keywords (e.g., “toxin,” “venom,” “knottin,” “kunitz,” “defensin”) detected in annotation descriptions from UniProt and eggNOG.

Based on these evidence types, peptides were classified into three categories according to annotation evidence and secretion support. First, identify high-confidence toxins that have at least one line of strong toxin-specific evidence and a positive secretion prediction. Second, toxin-like peptides with strong toxin-specific evidence but lacking secretion support. This was included because some true signal peptides may not be detected leading to a false negative. Third, all remaining peptides were classified as unlikely toxins.

To compare expression patterns among toxin annotation categories, analyses were restricted to the subset of transcripts corresponding to the top 10% most highly expressed transcripts per species. Within each sample, expression distributions of high-confidence toxins and toxin-like peptides were compared against unlikely toxins using two-sided Wilcoxon rank-sum tests. Resulting p-values were adjusted for multiple testing using the Benjamini–Hochberg procedure, and adjusted p-values (Padj) were used to assess statistical significance.

Identified toxin candidates were further categorized into broad functional categories based on sequence similarity to known toxin types. Peptides were assigned to the inhibitor cystine knot (ICK) category if supported by knottin-specific database matches or characteristic cysteine-rich motifs. Enzymatic toxins were defined based on annotations indicating catalytic activity, including proteases, phospholipases, and related enzyme classes. Other venom-associated peptides include sequences with annotation-supported venom relevance that did not fall into major structural or enzymatic toxin classes, whereas unclassified venom components represent peptides with evidence of secretion and toxin-like characteristics but without informative functional annotation. Toxin candidates were assigned to a single functional class and were summarized using counts of transcripts.

To further resolve venom composition, toxin family assignments were obtained based on similarity to curated UniProt toxin families. For each sample, toxin family composition was quantified by counting occurrences of annotated families, allowing transcripts with multiple annotations to contribute to multiple family counts. To enable comparison across samples, the most abundant toxin families per sample were identified and combined into a union set, with less frequent families grouped into an “Other families” category. Toxin family composition was visualized as stacked bar plots of occurrence counts across samples.

To further examine sequence-level conservation among highly expressed ICK peptides, peptides containing ICK motifs were selected from these transcripts based on prior functional classification. To enable comparative analysis of sequence features, peptide sequences were aligned using MAFFT v7.526 with the L-INS-i algorithm and 1,000 iterations of iterative refinement [28], and poorly aligned or gap-rich columns were removed using trimAl v1.5.1 [29]. A maximum likelihood phylogenetic tree was reconstructed using IQ-TREE v3.0.1 [30] with automatic model selection via ModelFinder (-m MFP), 1,000 replicates of ultrafast bootstrap approximation (-bb 1000), and 1,000 replicates of SH-aLRT branch support (-alrt 1000). The resulting tree was visualized as a cladogram alongside the multiple sequence alignment using R packages ggtree v4.0.4 [31] and ggplot2 [32]. Predicted signal and mature peptide regions were inferred based on peptide cleavage predictions derived from the ProP and SignalP predictions included in the consolidated annotation table described above. Conserved cysteine residues characteristic of ICK peptides are indicated in the alignment as gold-highlighted columns corresponding to the six canonical ICK disulfide-forming cysteines (C1-C6).

## Results

### Species identification

Species identification was carried out using established taxonomic standards for the subfamily *Selenocosmiinae*. Morphological analysis of the spider specimens revealed distinct characteristics which enabled species discrimination and classification under known genus and/or family. *Orphnaecus sp. ‘Mindanao 1’, Orphnaecus kwebaburdeos, Orphnaecus sp. L2, and Orphnaecus sp.* showed characteristics consistent to that of *Orphnaecus* genus, including having a reinform patch lyrate. *Selenobrachys philippinus* and *Selenobrachys ustromsupasius* showed characteristics congruent to that of the described *Selenobrachys* genus (Acuña, et al, 2025), including ovoid proximally truncated lyrate patch and tombstone-shaped spermathecal lobe. *Selenocosmia* gen. ‘A’ *et sp.* 1 and *Selenocosmia* gen. ‘A’ *et sp*. 2 showed possibility of a new genus, highlighting characteristics of a *Selenocosmiinae* subfamily but noncongruent to that of *Orphnaecus, Selenobrachys, Phlogiellus*.

### Transcriptome assembly and annotation

In this study, we have sequenced the transcriptomes of nine (9) venom glands, two (2) bodies, and one (1) each of chelicerae, abdomen, and leg, across eight (8) species within three (3) genera.. After basecalling, adapter trimming, and filtering, long-read sequencing of venom gland tissues generated a total of 62M high quality reads across all specimens (min. 0.8M, max 7.9M reads per sample), with a mean read length of 625 bp (Table 1). *De novo* assembly yielded between 2,318 and 12,923 transcripts per sample, from which 2,864 to 13,014 candidate peptide sequences were predicted, potentially reflecting the recovery of isoforms and multiple ORFs from some transcripts (Table 1). Across all samples, 65% of predicted peptides were successfully annotated using eggNOG, indicating substantial functional annotation coverage. Collectively, the sequencing depth provided sufficient coverage to recover thousands of predicted peptide sequences per sample, particularly highly expressed toxin-associated transcripts, which formed the basis of the comparative analyses presented in this study.

**Table 1.**
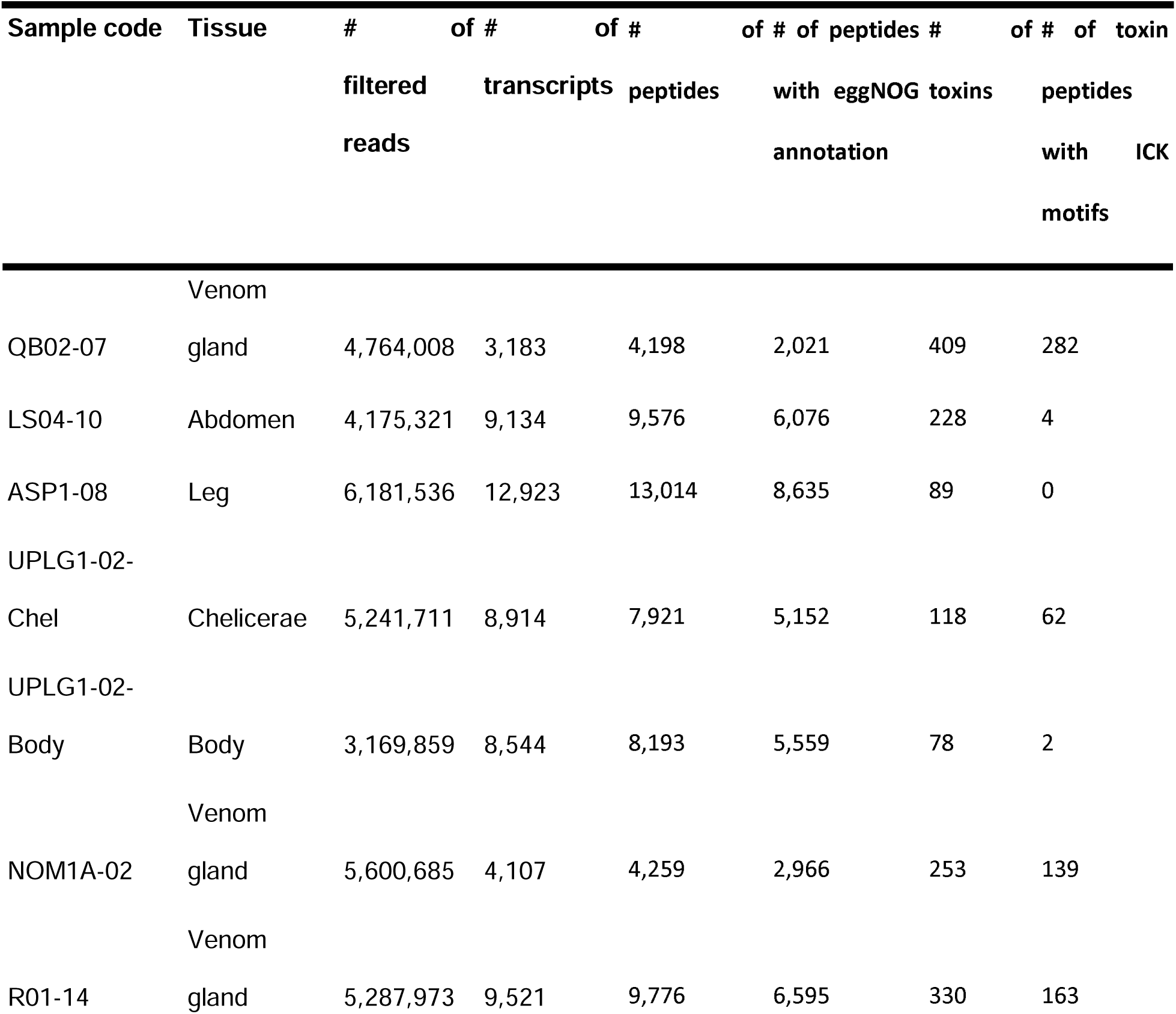
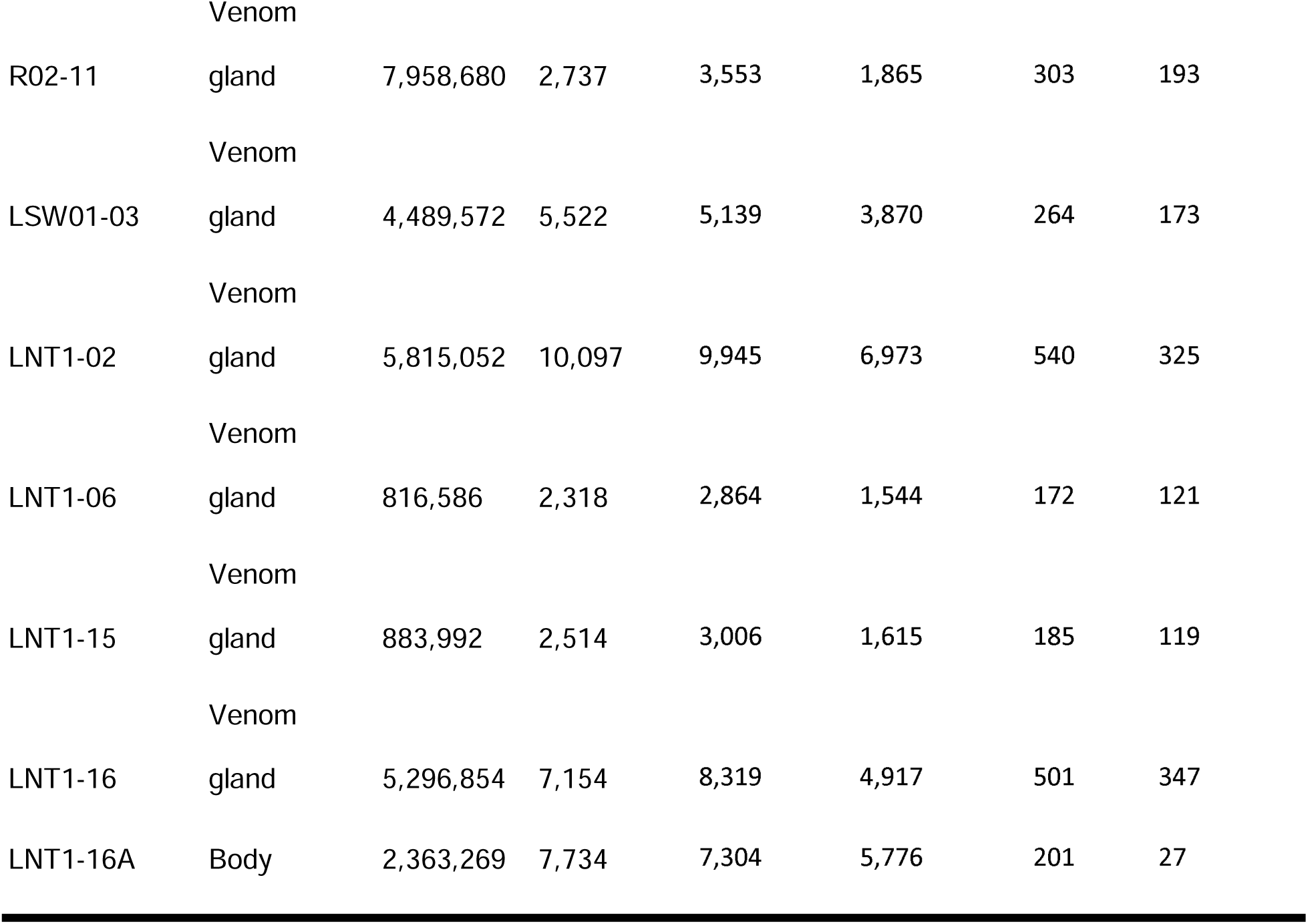
Transcriptome assembly metrics, functional annotation statistics, and predicted toxin content of venom gland and non-venom tissue transcriptomes.

Venom gland transcriptomes consistently exhibited a higher proportion of toxin-associated peptides compared to non-venom tissues. The number of predicted toxins ranged from 118 to 540 in venom gland and chelicerae samples, compared with the substantially lower range of 78 to 228 in non-venom tissues (abdomen, leg, and body) despite some non-venom samples having comparable or higher numbers of transcripts and predicted peptides. Similarly, peptides containing inhibitor cystine knot (ICK) motifs were strongly enriched in venom gland transcriptomes, ranging from 62 to 347 peptides whereas non-venom tissues contained only 0 to 27 ICK-containing toxin peptides. Interestingly, some non-venom tissues exhibited higher total transcript and peptide counts than venom glands, possibly reflecting broader baseline gene expression. However, venom glands generally showed greater toxin representation, consistent with more targeted expression of toxin-rich gene families.

### Phylogenetic context and orthogroup distribution

To provide phylogenetic context for venom transcriptome comparisons, a species tree was inferred using OrthoFinder from predicted peptide sequences across all sampled specimens. OrthoFinder clustered homologous peptides into orthogroups, reconstructed gene trees for individual orthogroups, and used these gene trees to infer a species phylogeny. The resulting species phylogeny was congruent with established relationships among theraphosid spiders, with sampled taxa clustering according to their expected taxonomic affiliations (Figure 2A). Three distinct monophyletic clades were observed: *Selenobrachys philippinus* (NOM1A-02) and *Selenobrachys ustromsupasius* (R02-11 & R01-14) formed one clade (pink in Fig. 2a), *Orphnaecus kwebaburdeos* (QB2-07), *Orphnaecus* sp. M1 (ASP01-08), *Orphnaecus* sp. L2 (LS04) and *Orphnaecus* sp. (UPLG1) formed another clade (light blue in Fig. 2a), and taxa from the Selenocosmiinae genus A (LNT1 and LSW01) formed a separate lineage from the other clades (light green in Fig. 2A), which is consistent with the known phylogenetic relationships of the sampled species.

**Figure 1.**
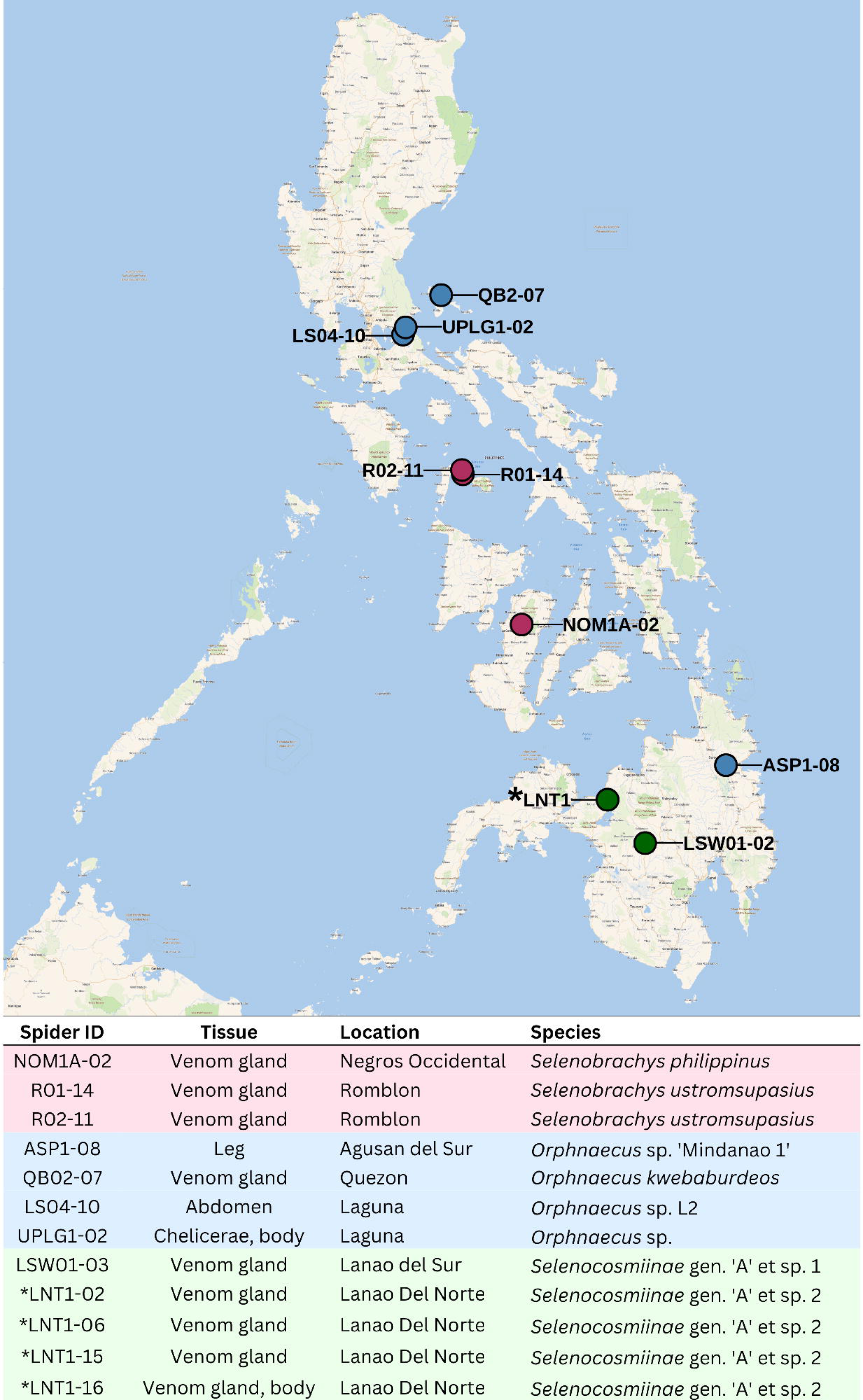
Sampling locations and transcriptome dataset overview. Sampling localities of Philippine tarantulas included in this study are shown on the map, with colors corresponding to genus. The accompanying table summarizes the transcriptome datasets, including sample ID, tissue source, collection locality, and taxonomic assignment. Asterisks indicate multiple samples from the same location.

**Figure 2.**
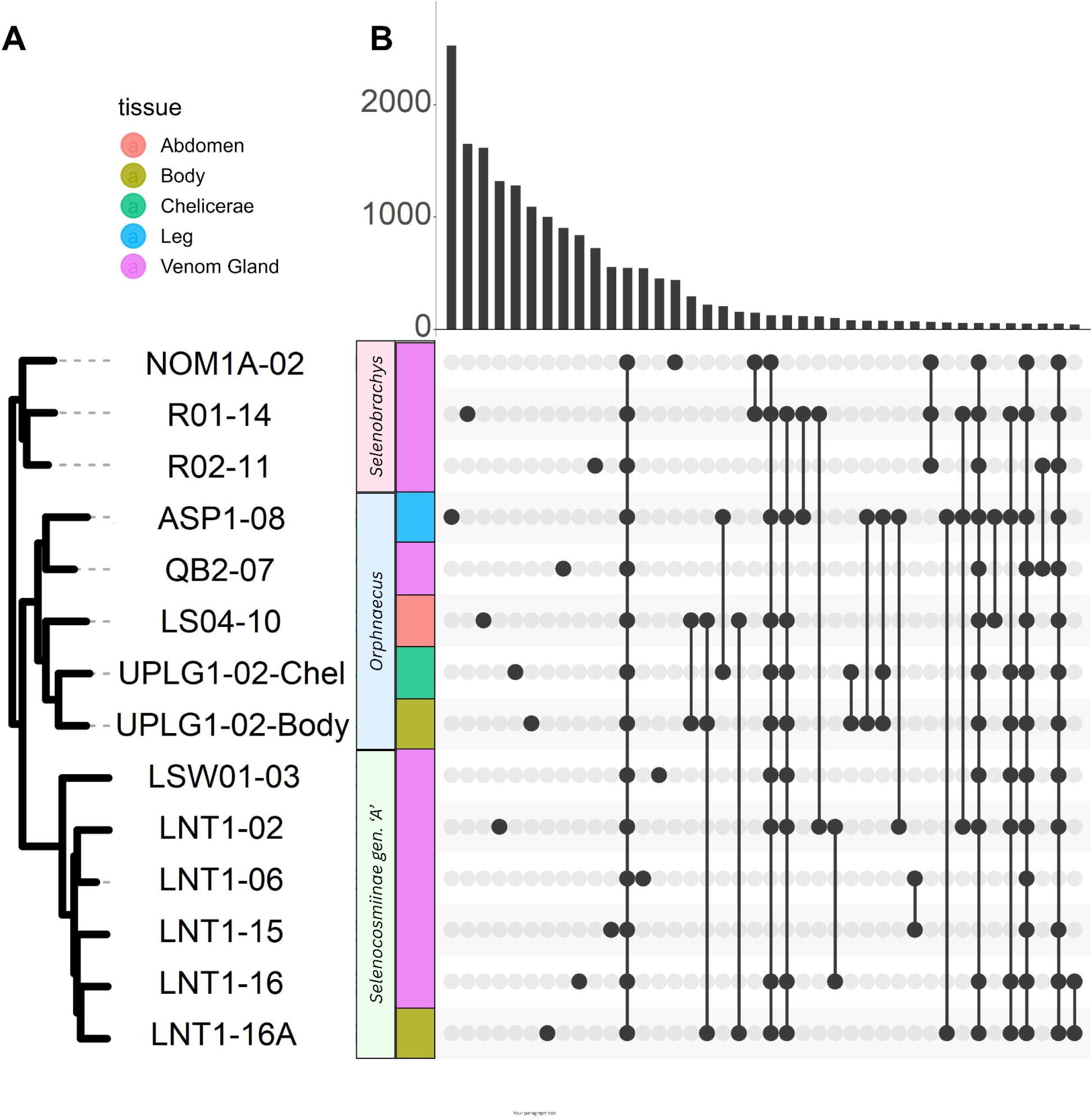
Orthogroup distribution and phylogenetic relationships of sampled taxa. (A) Species tree inferred from orthologous genes recovered from all sampled transcriptomes. Clades are labelled according to genus, and tissue type is indicated by the adjacent color strip. (B) UpSet plot showing the distribution of orthogroups across transcriptomes. Vertical bars indicate the number of orthogroups detected in each unique combination of transcriptomes, whereas connected dots identify the transcriptomes contributing to each intersection. The plot illustrates both broadly shared orthogroups and orthogroups restricted to specific genera, species, or individual transcriptomes.

Clustering of predicted peptides using OrthoFinder grouped homologous peptides into orthogroups, enabling comparison of gene families across samples. Analysis of orthogroup distribution revealed both shared and lineage-specific orthogroups (Figure 2B). In the UpSet plot, each bar represents the number of orthogroups shared among a particular combination of samples. A substantial proportion of orthogroups were present across multiple species, indicating conserved peptide families, while others demonstrate sample, tissue, and lineage specificity, reflecting both biological specialization and lineage-specific diversification.

### Venom gland specialization

Principal component analysis (PCA) of orthogroup-level expression profiles revealed separation of samples according to tissue type along the first principal component (PC1) (Figure 3A). PC1 explained 21.7% of the total variance, while PC2 accounted for 15.8%. Venom gland samples formed a distinct cluster along the positive direction of PC1, whereas body, abdomen, leg, and cheliceral tissues grouped along the negative axis, indicating a shared non-venom transcriptional background. This separation was statistically supported, with tissue identity significantly explaining variation along PC1 (ANOVA, F = 18.964, P = 0.0002), consistent with a strong tissue-driven transcriptional axis conserved across species.

**Figure 3.**
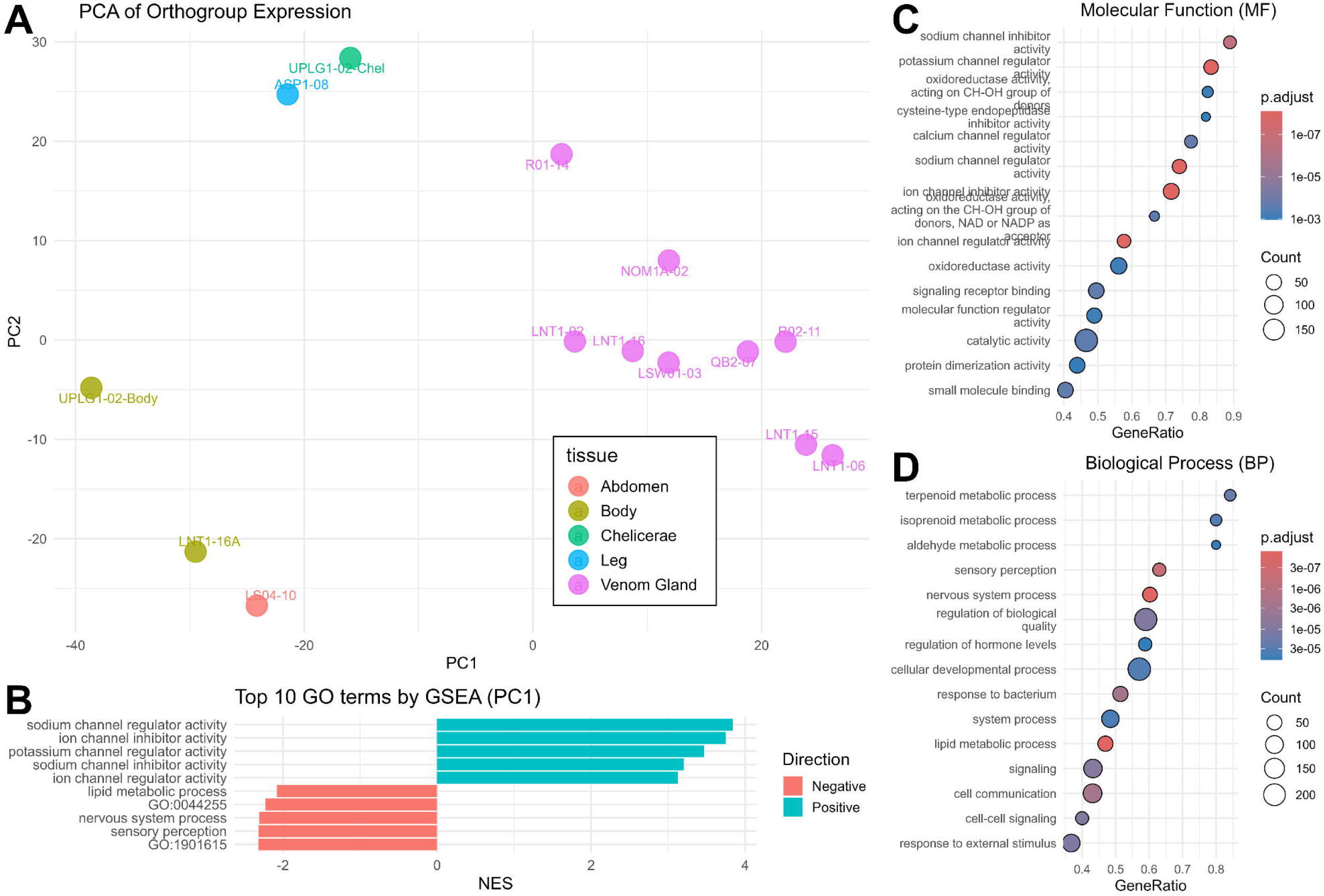
Venom gland transcriptomes exhibit distinct orthogroup expression profiles enriched for toxin-associated functions. (A) Principal component analysis (PCA) of orthogroup expression profiles based on the top 20% most variable orthogroups. Samples are colored according to tissue type (B) Gene Set Enrichment Analysis (GSEA) of orthogroups ranked according to their loadings on PC1, showing the top positively and negatively enriched Gene Ontology (GO) terms. Positive normalized enrichment scores (NES) correspond to orthogroups associated with venom gland samples, whereas negative NES correspond to orthogroups associated with non-venom tissues. (C–D) Dot plots showing significantly enriched GO Molecular Function (MF) and Biological Process (BP) terms identified by GSEA. GeneRatio represents the proportion of orthogroups associated with each GO term. Dot size indicates the number of orthogroups associated with each GO term, and color denotes the adjusted P-value.

To identify the molecular drivers underlying this separation, orthogroups were ranked based on their loadings on PC1. Examination of PC1 loadings indicated that the observed separation of tissues on PC1 was driven by a coordinated set of orthogroups rather than a single dominant feature. Among the top contributors to PC1, orthogroups with positive loadings, which are associated with venom gland samples, include functions related to ion channel regulation and inhibition, consistent with venom-associated neurotoxic activity (Suppl Fig 2). Gene Set Enrichment Analysis (GSEA) corroborated this trend, emphasizing the enrichment of ion channel regulatory and inhibitory activities (Figure 3B), alongside broader molecular functions such as binding and catalytic processes (Figure 3C–D).

The consistent separation of venom gland samples and the enrichment of toxin-associated functions relative to non-venom tissue samples support a conserved transcriptional signature underlying venom gland specialization. These findings suggest that venom gland specialization extends beyond toxin gene expression and involves broader cellular processes associated with venom gland function. Together, this functional partitioning defines a distinct molecular signature of venom gland transcriptomes across theraphosid spiders.

### Toxin composition and diversity

Classification of predicted peptides into high-confidence toxins, toxin-like peptides, or non-toxins based on sequence similarity and annotation criteria revealed substantial variation in toxin representation across tissues (Figure 4A). Within-sample expression levels of toxin-associated transcripts were significantly higher than non-toxin transcripts across venom gland and chelicerae samples (Wilcoxon rank-sum test with Benjamini–Hochberg correction; Padj < 0.05 to Padj < 0.0001). This pattern was consistent across all species, indicating a conserved enrichment of toxin-associated transcripts in venom-associated tissues.

**Figure 4.**
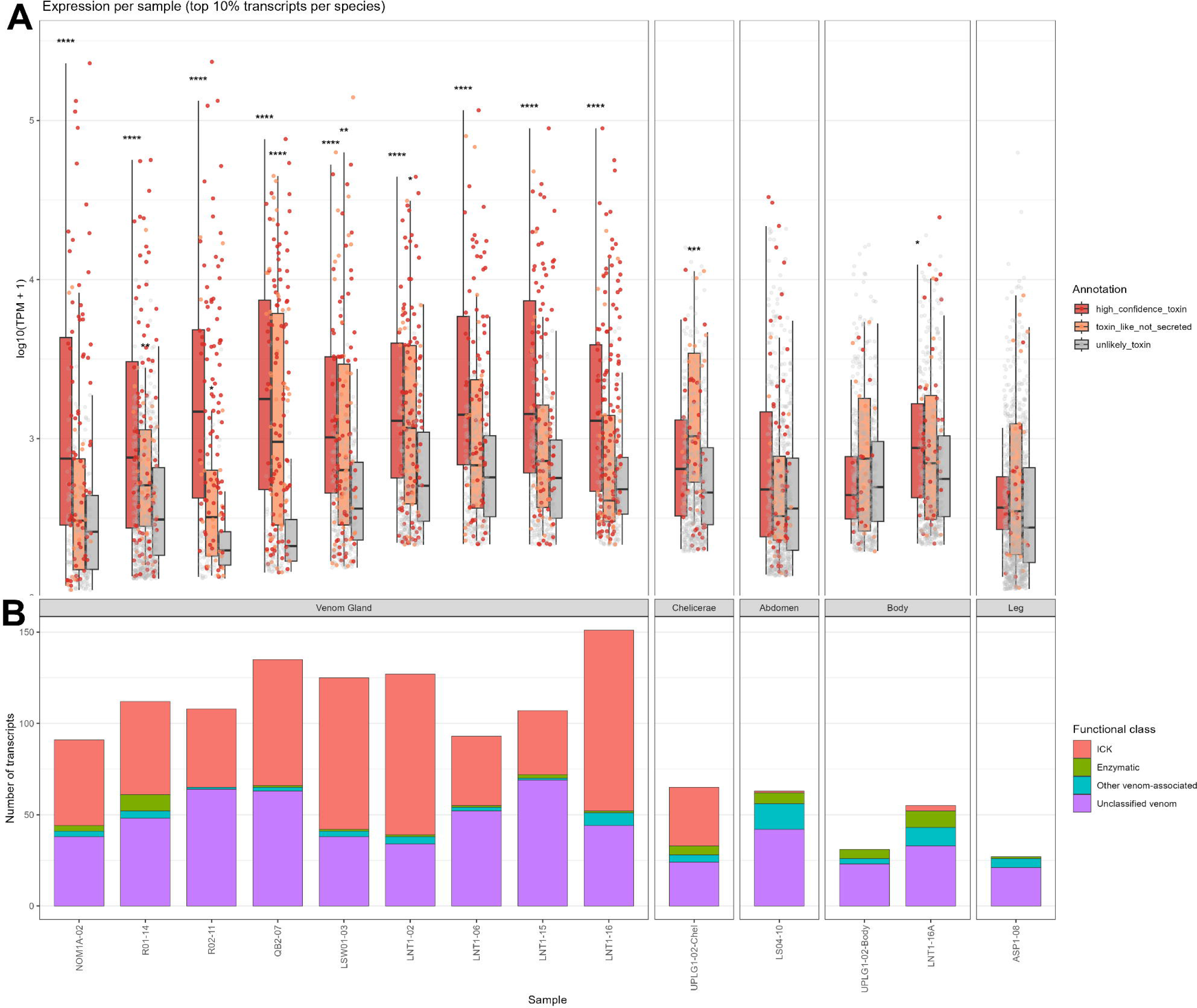
Venom gland transcriptomes contain highly expressed toxin transcripts, including abundant inhibitor cystine knot peptides. (A) Distribution of transcript abundance (log_10_(TPM+1)) among the top 10% most highly expressed transcripts in each sample, grouped according to toxin annotation category. Boxplots summarize the distribution of transcript abundance, with points representing individual transcripts. Statistical significance indicates comparison with “unlikely toxin” category after multiple-testing correction. (B) Functional composition of the top 10% most highly expressed toxin transcripts across samples showing the number of transcripts assigned to each toxin functional category.

Functional categorization of toxin candidates further revealed that cysteine-rich peptide toxins represented the largest fraction of venom-associated transcripts, with those encoding ICK peptides accounting for approximately 50% of toxin-classified transcripts in venom glands (Figure 4B). Enzymatic toxins, including proteases and phospholipase-related proteins, as well as other venom-associated peptides, were also consistently represented, although at lower proportions. Notably, ICK peptides were strongly enriched in venom gland samples and were largely absent or minimally represented in non-venom tissues, consistent with their role as core components of spider venom. Interestingly, a substantial number of ICK-encoding transcripts were also detected in cheliceral tissues, indicating that these structures may serve as an alternative source for toxin studies when intact venom glands are not accessible, particularly in smaller specimens. Together, these results highlight the dominance of cysteine-rich peptide toxins within Philippine tarantula venom gland transcriptomes.

Toxin family annotation revealed substantial diversity within venom gland transcriptomes across all specimens (Figure 5). Predicted toxin peptides were assigned to multiple toxin families based on sequence similarity and structural features, with individual samples containing a broad repertoire of families. Several toxin families were shared across specimens, indicating a conserved core venom repertoire, while additional families were restricted to a subset of samples, suggesting lineage- or specimen-specific diversification. Among these, the neurotoxin 10 family was consistently the most abundant and widely represented across all venom gland samples. The neurotoxin-14 family was also detected across specimens, although in lower abundance. Additional families, including neurotoxin 3 and subfamilies 02, 07, and 34, were present in nearly all venom gland samples but showed reduced representation particularly in NOM1A-02.

**Figure 5.**
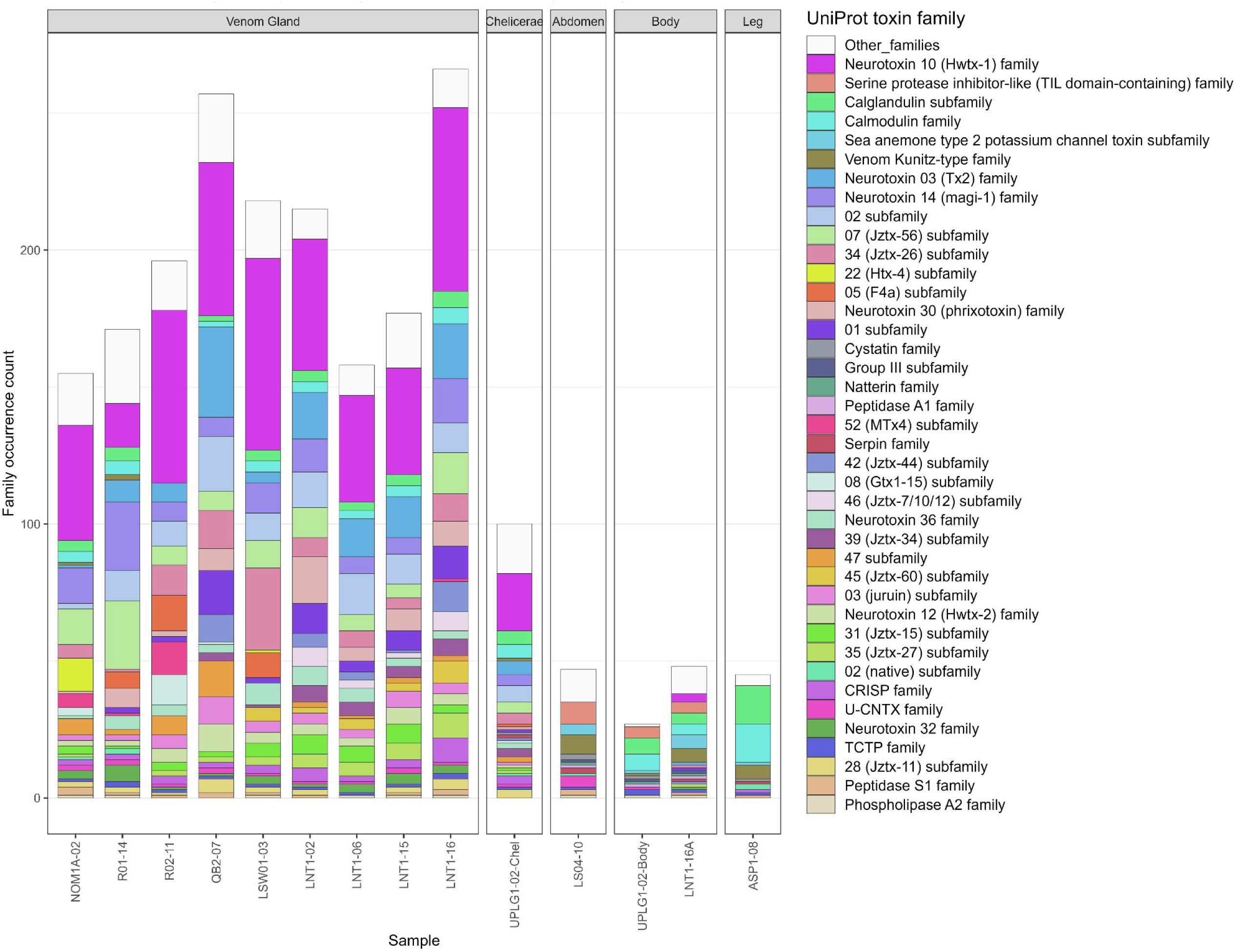
Composition and diversity of UniProt toxin families. Stacked bar plots showing the number of predicted toxin peptides assigned to each UniProt toxin family across transcriptomes. Colors represent individual toxin families, whereas white segments denote all remaining toxin families not individually displayed in the legend.

In contrast, calcium-binding proteins such as calglandulin and calmodulin were detected across most samples and were not restricted to venom glands. These proteins were absent in the R01-14 venom gland and LS04-10 abdomen samples, and were most abundant in leg tissue, suggesting that these proteins primarily serve conserved physiological functions rather than venom-specific roles. While a subset of toxin families contributed substantially to overall venom composition, many additional families were detected at lower frequencies and were grouped into an “Other families” category.

Overall, these results indicate that venom gland transcriptomes are characterized by both a conserved core of dominant venom-associated families and a diverse repertoire of lower-abundance components, suggesting that conserved venom-associated families likely support core venom functions, while lineage-specific families contribute to venom diversification.

### Sequence conservation and diversification of dominant ICK peptides

Since ICK-type toxins were identified as a dominant component of venom gland transcriptomes, further analyses were performed on these peptides. To examine sequence-level conservation, the highest-expressed transcripts per sample were identified, and their corresponding predicted peptide sequences were selected for alignment and analysis. Notably, all top transcripts were classified as ICK-type toxins (Figure 6). All sequences exhibited a conserved cysteine framework consistent with the canonical ICK motif, with six cysteine residues (C1–C6) maintained across samples. Full-length transcript sequences, spanning from the 5’ cap to the 3’ poly(A) tail, and near full-length transcripts were assembled, which enabled clear identification of precursor structure, including an N-terminal Signal peptide, followed by variable propeptide regions and a cysteine-rich mature peptide domain.

**Figure 6.**
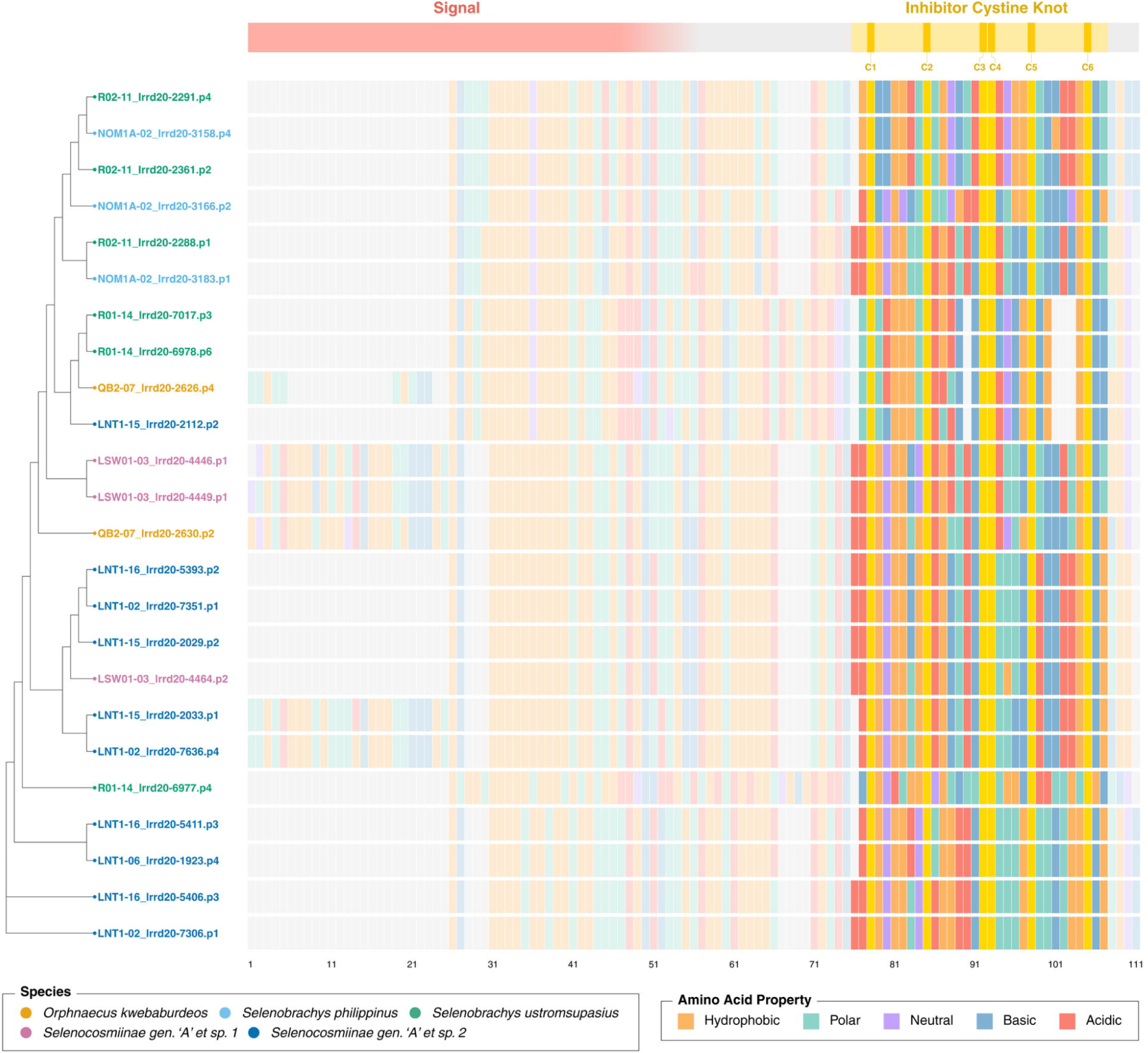
Highly expressed inhibitor cystine knot peptide precursors retain a conserved cysteine framework. Multiple sequence alignment and phylogenetics tree of the highest-expressed inhibitor cystine knot (ICK) peptide precursors identified from venom gland transcriptomes. The signal peptide, propeptide, and mature peptide regions are indicated above the alignment. Conserved cysteine residues (C1–C6) defining the canonical ICK framework are highlighted, whereas variation is observed primarily within the inter-cysteine regions of the mature peptide. Amino acids are colored according to their physicochemical properties.

Despite this conserved structural architecture, substantial sequence variation was observed in the inter-cysteine regions across samples. Phylogenetic analysis based on aligned peptide sequences revealed the clustering of ICK peptides according to sequence similarity, with multiple distinct groups observed across samples. While some clusters were consistent with sample identity, others reflected shared sequence features across species, indicating both lineage-specific diversification and conservation of toxin scaffolds.

## Discussion

This study presents the first long-read transcriptomic analysis for Philippine tarantulas that focuses on venom glands, and provides insights into toxin precursor diversity and venom gland specialization. By leveraging long-read sequencing, we were able to recover full-length and near full-length toxin precursor transcripts, enabling characterization of signal peptides, propeptide regions, and mature toxin domains. This represents an important advantage for venom studies, where highly similar toxin genes and repetitive sequence motifs can complicate transcript reconstruction and precursor annotation in short-read datasets.

We identified peptides representing diverse functional classes, including non toxin components, while the majority of predicted peptides were classified as toxins, specifically those involved in sodium channel regulation. We further examined 136 cysteine rich peptides from three species representing three genera (*Orphnaecus, Selenobrachys*, and an undescribed genus from the Selenocosmiinae subfamily). Phylogenetic analysis of toxin peptides revealed distinct inhibitory cysteine knot (ICK) families which could represent potentially novel ICK families. Interestingly, some of these peptides may be either shared or unique across species indicating lineage specificity. Indeed, the distribution of these peptide families across different species is not equal, suggesting a complex process in the evolution of these ICK families in the tarantula venom gland.

These findings underscore the complexity of tarantula venoms and their potential applications in drug discovery and biotechnology. By elucidating the diversity of venom peptides, this research contributes to our understanding of venom evolution and the biological functions of these compounds.

### Phylogeny and orthology

The orthogroup-based phylogenetic reconstruction recovered relationships that were largely congruent with the current taxonomic placement of the sampled theraphosid spiders. Specimens belonging to the genera Selenobrachys and Orphnaecus formed distinct clades, while the sampled representatives of the putative Selenocosmiinae genus A formed a separate lineage. This agreement between transcriptome-derived phylogenetic inference and morphology-based taxonomy supports the utility of venom gland transcriptomic datasets for resolving evolutionary relationships among closely related theraphosid taxa. In particular, the distinct placement of the Selenocosmiinae genus A lineage provides additional molecular evidence supporting its separation from currently recognized genera, although broader taxon sampling will be required to evaluate its formal taxonomic status.

Orthogroup clustering further revealed a combination of conserved and lineage-specific gene families. A substantial proportion of orthogroups were shared across multiple species, indicating the presence of a conserved set of genes maintained across theraphosid lineages. Such conserved orthogroups likely represent fundamental biological functions, including cellular processes associated with venom production, secretion, and maintenance. At the same time, the presence of lineage-specific orthogroups suggests ongoing evolutionary diversification following lineage divergence. Similar patterns have been reported across venomous animals, where gene duplication, sequence divergence, and differential retention contribute to the expansion of lineage-specific venom gene repertoires [9,33]

The coexistence of shared and lineage-specific orthogroups supports a model in which venom systems evolve through the interaction of evolutionary conservation and innovation. Conserved gene families provide the molecular foundation for venom gland function, whereas lineage-specific genes may contribute to species-level differences in venom composition and ecological adaptation. Although the present study was not designed to investigate the evolutionary mechanisms underlying orthogroup diversification, the observed patterns are consistent with previous studies showing that venom repertoires are shaped by both phylogenetic inheritance and lineage-specific diversification of toxin-associated gene families [34,35].

### Venom gland specialization

The observed separation of venom gland and non-venom tissues in the PCA supports the existence of a conserved transcriptional profile associated with venom gland specialization across theraphosid spiders. Similarly, the predominance of non-toxin transcripts in non-venom tissues further supports the functional partitioning of gene expression across tissue types [36]. This supports the presence of a conserved transcriptional framework associated with venom gland function [37].

While toxin genes are often assumed to drive this separation due to their high expression and tissue specificity, the coordinated contribution of multiple orthogroups in our analysis indicates that this pattern is not dictated by a single gene class. Instead, the observed separation reflects broader differences in gene expression associated with venom gland specialization, suggesting that venom gland function depends on a diverse set of genes beyond toxin-encoding transcripts. Indeed, toxin production is supported by complex cellular and regulatory processes, including cell-type-specific expression patterns and regulatory mechanisms that coordinate toxin synthesis and secretion [37,38].

In addition, venom gland transcriptomes in diverse venomous animals are enriched for genes involved in regulation, signaling, and metabolic processes in addition to toxin production [37]. The enrichment patterns observed in this study are consistent with this trend, suggesting that venom gland specialization extends beyond toxin expression and involves the molecular machinery required for toxin synthesis, processing, regulation, and deployment. Together, these findings support the emerging view that independently evolved venom glands share conserved transcriptional programs associated with protein synthesis, secretion, and cellular regulation, despite substantial lineage-specific differences in toxin repertoires [37,39] . The transcriptional patterns observed in Philippine theraphosid venom glands are compatible with this broader framework, suggesting that similar molecular processes may underlie venom gland specialization in this understudied lineage.

### Toxin composition and diversity

Toxin family annotation revealed a complex venom composition characterized by both conserved and variable components. The presence of widely shared toxin families, including highly abundant neurotoxin 10 and the consistently detected neurotoxin 14, supports the consistent conserved core venom repertoire across Philippine tarantulas.

While several toxin families were shared across specimens, additional families were detected in a subset of specimens, suggesting lineage- or specimen-specific diversification. Indeed, comparative venomic analyses have consistently reported such patterns of conserved toxin combined with lineage-specific toxin expansions, showing both functional constraints and rapid evolutionary diversification of venom gene families [35].

The enrichment of high-confidence toxins and toxin-like peptides in venom gland transcriptomes relative to non-venom tissues observed in this study is consistent with the specialized secretory function of venom glands. Such tissue-specific enrichment of toxin transcripts has been widely reported in spider venomics studies, where venom glands exhibit distinct expression profiles compared to other body tissues [40].

The dominance of cysteine-rich peptide toxins, particularly ICK peptides, in venom gland samples highlights their central role in tarantula venoms. Disulfide-rich peptides adopting the inhibitor cystine knot scaffold are well established as the principal structural and functional components of spider venoms, particularly in mygalomorph spiders such as Theraphosidae [41]. Their strong enrichment in venom glands and near-absence in non-venom tissues underscores their specialization for venom-related functions, particularly in modulating ion channels involved in prey immobilization and defense [42].

The detection of abundant ICK-encoding transcripts in cheliceral samples is consistent with the anatomical location of the venom gland within the chelicera. Notably, toxin-associated transcripts remained readily detectable despite the presence of additional cheliceral tissues, suggesting that cheliceral samples may represent a practical alternative for toxin discovery particularly in cases where venom glands are difficult to isolate.

Aside from toxin peptides, spider venoms frequently contain a diverse array of enzymatic and auxiliary proteins including metalloproteases, hyaluronidases, and cysteine-rich secretory proteins, which can facilitate toxic diffusion, tissue penetration, or synergistic venom activity [43]. In fact, proteomic surveys across Theraphosidae species revealed complex venom compositions consisting of both neurotoxic peptides and enzymatic proteins, underscoring the biochemical diversity of tarantula venoms [44]. Consistent with this complexity, calcium-binding proteins such as calglandulin and calmodulin were detected across multiple tissue types in this study. Their widespread distribution beyond venom glands suggests broader physiological functions rather than venom-specific roles, although related proteins have been implicated in venom secretion and modulation of venom activity in other venomous animals [45]. Together, these findings highlight the dynamic and complex nature of venom systems, characterized by a core of functionally important toxins alongside a diverse array of auxiliary and lineage-specific components.

### Inhibitory cysteine knot peptides

Inhibitor cystine knot ICK peptides comprise the largest and most functionally significant toxin class in spider venoms [35,46]. The predominance of ICK peptides in Philippine tarantula transcriptomes reinforces their central role as the primary functional components of theraphosid venoms. The consistent recovery of the ICK toxins as the most highly expressed transcripts across all venom gland samples highlights their conserved biological importance, likely reflecting strong selective pressures to maintain their role in prey immobilization and defense.

Despite the conserved cysteine framework defining the ICK scaffold, extensive variation in inter-cysteine regions was observed supporting the hypothesis that functional diversification is driven primarily by sequence variation outside disulfide-bonded cores. This pattern aligns with established models of toxin evolution, where structural stability is maintained by conserved cysteine residues while adaptive diversification occurs through mutations in functionally exposed regions [47,48]. These results support a model in which the evolution of theraphosid venom is driven by the preservation of a conserved structural scaffold and the rapid diversification of functional residues.

Phylogenetic clustering revealed a dual pattern of diversification. While some ICK peptides grouped according to species identity, others clustered across taxa, suggesting coexistence of lineage-specific expansions and conserved toxin lineages. Indeed, such patterns are consistent with evolutionary processes including gene duplication, divergence, and potential convergence in venom gene families [9,46,49].

The high diversity of ICK peptides identified in the study supports previous reports describing this toxin class as the most abundant and variable component of spider venoms [4,46] (Saez et al., 2010, Pined et al., 2020). A key strength of this study lies in the use of full-length and near full length transcript sequences. Unlike short-read assemblies, which often fragment toxin precursors or obscure gene structure, this approach enabled accurate reconstruction of complete toxin precursor architecture, including signal peptides, propeptide regions, and mature toxin domains [33]. To our knowledge, this represents the first application of long-read transcriptomics to characterize venom gland peptide diversity in Philippine Theraphosidae, providing a structurally resolved view of toxin repertoires and a foundation for future studies of venom evolution and functional diversification.

## Conclusion

Together, the results in this paper suggest that theraphosid venom evolution is characterized by conservation of venom gland transcriptional architecture and ICK structural scaffolds, coupled with diversification of toxin repertoires and functional residues. However, transcriptomic data alone cannot fully resolve the genomic mechanisms underlying toxin repertoire expansion. Future whole-genome sequencing efforts will be essential for identifying gene duplication events, genomic organization of toxin loci, and other evolutionary processes that contribute to venom diversification.

While this study provides a comprehensive transcriptomic resource for Philippine theraphosid venom glands, further complementary approaches will expand the biological insights gained from these data. Toxin annotation was based on transcriptomic evidence and sequence homology, providing a robust framework for candidate toxin discovery that can be further prioritized through structural modeling and molecular dynamics simulations, and validated through functional in vitro and in vivo assays. Likewise, transcript abundance reflects gene expression within the venom gland but may not directly correspond to the abundance of mature peptides in secreted venom. Lastly, expanding taxonomic sampling across additional Philippine theraphosid lineages will further improve our understanding of venom diversity and evolution. Nevertheless, the integration of long-read transcriptomics in this study establishes a robust molecular resource for future investigations into toxin evolution, functional characterization, and the biotechnological potential of theraphosid venom peptides.

## Acknowledgments

The authors would like to extend its gratitude to the Department of Science and Technology (DOST) for funding the research along with the National Research Council of the Philippines (NRCP). The authors likewise express their sincere appreciation to the Department of Environment and Natural Resources–Biodiversity Management Bureau (DENR-BMB) for granting the permit (Gratuitous Permit No. 318) necessary for the conduct of sampling activities, and to the local government units and concerned authorities in Brgy. Tambac and Brgy. Guimpingan, Municipality of Romblon, Romblon Island; Brgy. Poblacion, Municipality of Prosperidad, Agusan del Sur; Brgy. Aluyon, Burdeos, Municipality of Quezon Province, Luzon Island; Brgy. Halayhayin-Balian, Municipality of Siniloan, Province of Laguna, Luzon Island; UP Sierra Madre Land Grant, Laguna-Quezon, Luzon Island; Brgy. Amoyong, Municipality of Wao, Lanao del Sur; and Municipality of Tagoloan, Lanao Del Norte.

## Author’s contributions

L.R.P.R., L.A.G., and M.R.S.-B. conceived the study and managed the project. L.R.P.R. performed the experimental work, curated the data, conducted the formal analyses, prepared the figures, and drafted the manuscript. M.M.U.D. performed experiments, contributed to the formal analyses, and assisted in manuscript preparation. S.A.S.G. conducted the inhibitor cystine knot (ICK) peptide analyses. A.G.M.B. performed the phylogenetic analyses. D.C.A. performed venom gland dissections and maintained the spider collection. H.L.F.-C. and R.C.H.d.R. supervised the study. M.R.S.-B. and L.A.G. secured funding. All authors contributed to manuscript preparation, reviewed and edited the manuscript, and approved the final manuscript.

## Data availability

Raw sequencing data and transcriptome assemblies can be accessed at NCBI under BioProject accession number PRJNA1228174

