## Supplemental Figures for "Long-read transcriptomics highlights venom gland specialization and Inhibitor Cystine Knot (ICK) rich toxin diversity in Philippine tarantulas"

### Supplemental Figure 1

#### top 10% most variable OG

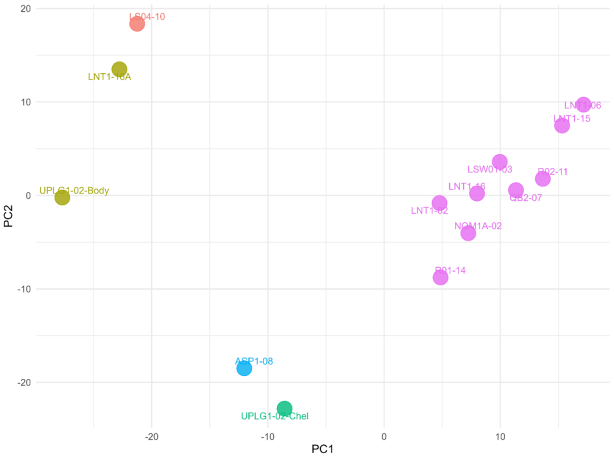


#### top 20% most variable OG

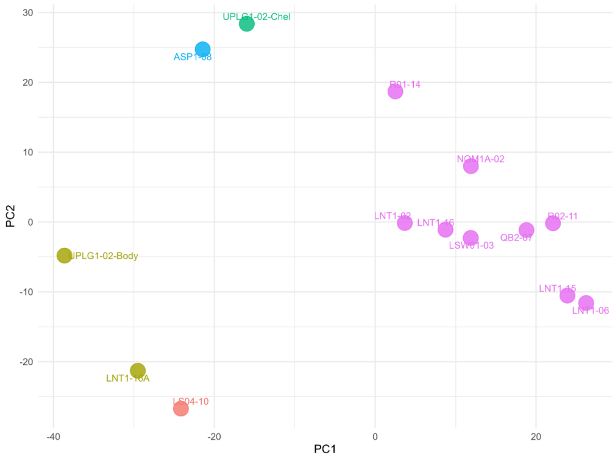


#### top 30% most variable OG

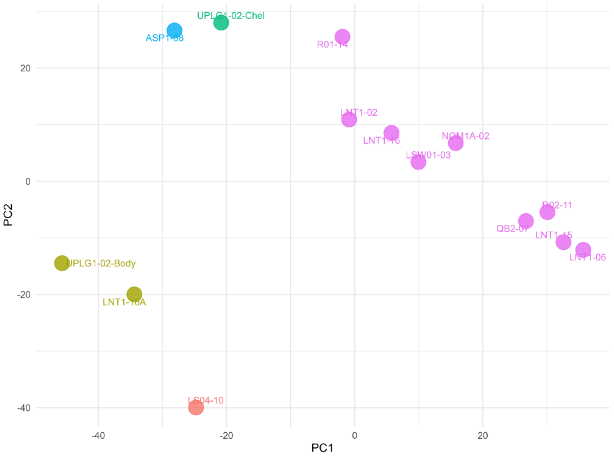


#### top 40% most variable OG

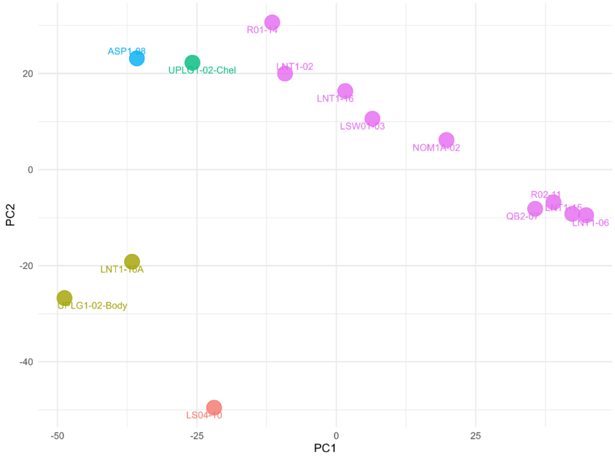

### Supplemental Figure 2

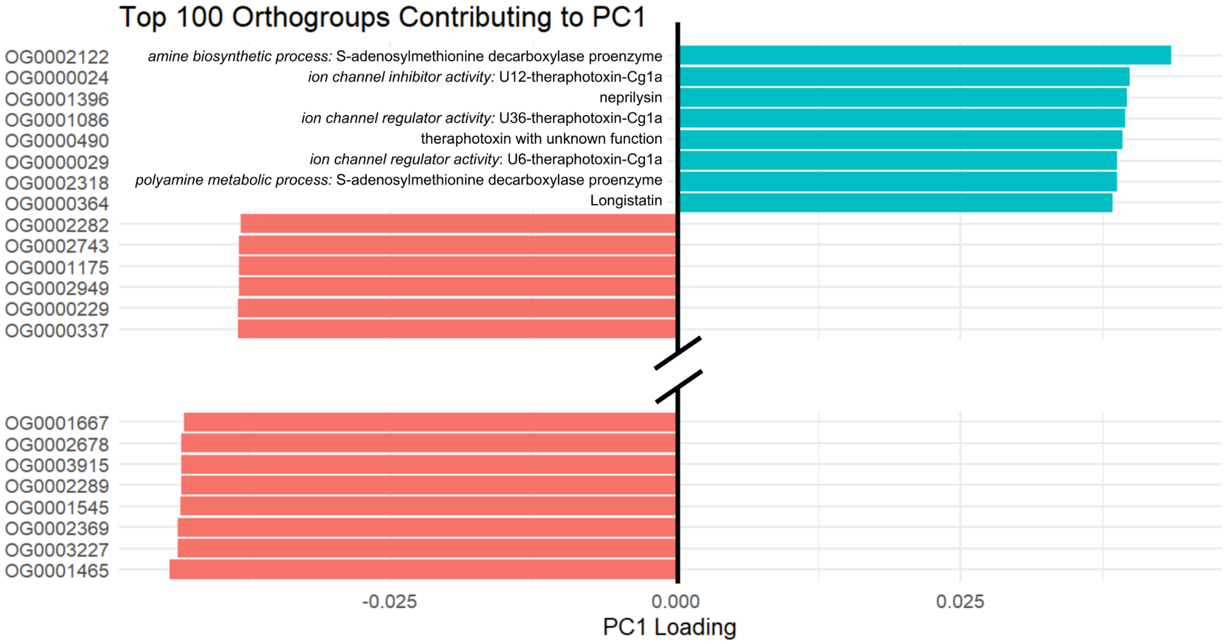
